# A catalog of single nucleotide changes distinguishing modern humans from archaic hominins

**DOI:** 10.1101/298950

**Authors:** Martin Kuhlwilm, Cedric Boeckx

## Abstract

Throughout the past decade, studying ancient genomes provided unique insights into human prehistory, and differences between modern humans and other branches like Neanderthals can enrich our understanding of the molecular basis of unique modern human traits. Modern human variation and the interactions between different hominin lineages are now well studied, making it reasonable to go beyond fixed changes and explore changes that are observed at high frequency in present-day humans. Here, we identify 571 genes with non-synonymous changes at high frequency. We suggest that molecular mechanisms in cell division and networks affecting cellular features of neurons were prominently modified by these changes. Complex phenotypes in brain growth trajectory and cognitive traits are likely influenced by these networks and other changes presented here. We propose that at least some of these changes contributed to uniquely human traits, and should be prioritized for experimental validation.

## Introduction

*Homo sapiens* appears to be a “very special primate” (Pääbo 2014). Our position among animal species stands out largely thanks to the composite complexity of our cultures, social structures and communication systems. It seems reasonable that this “human condition” is rooted, at least in part, in the properties of our brain, and that these can be traced to changes in the genome on the modern human lineage. This phenotype in the population called “anatomically modern humans” emerged in Africa likely before the deepest divergence less than 100,000-200,000 years ago (Schlebusch et al. 2012; Kuhlwilm et al. 2016), although complex population structure may reach back up to 300,000 years ago (Hublin et al. 2017; Schlebusch et al. 2017; Skoglund et al. 2017). Except for some early dispersals (Rabett 2018), humans most likely peopled other parts of the world than Africa and the Middle East permanently only after around 65,000 years ago. It has been claimed that the brain of modern humans adopted a specific, apomorphic growth trajectory early in life that gave rise to the skull shape difference between modern humans and extinct branches of the genus *Homo* (Hublin et al. 2015). Importantly, the growth pattern might differ between the populations (Gunz et al. 2010; Neubauer et al. 2018), while the brain size and encephalization of humans and Neanderthals is similar, with slightly larger brains in the latter (Trinkaus and Howells 1979; Schoenemann 2004; Hublin et al. 2015). This ontogenic trajectory, termed the “globularization phase”, might have contributed to cognitive changes that underlie behavioral traits in which humans differ from their extinct relatives, despite mounting evidence for their cognitive sophistication (Gunz et al. 2012; Hublin et al. 2015; Wynn et al. 2016; Boeckx 2017; Hoffmann et al. 2018).

We are now in a favorable position to examine the evolution of human biology with the help of the fossil record, in particular thanks to breakthroughs in paleogenomics: The recent reconstruction of the high quality genomes of members of archaic *Homo* populations (Meyer et al. 2012; Prüfer et al. 2014; Prüfer et al. 2017) has opened the door to new comparative genomic approaches and molecular analyses. The split of the lineages leading to modern humans and other archaic forms (Neanderthals and Denisovans) is estimated to around 600,000 years ago (Kuhlwilm et al. 2016), setting the timeframe for truly modern human-specific changes after this split, but before the divergence of modern human populations (Fig. 1). Together with efforts to explore present-day human diversity (Auton et al. 2015), this progress has allowed to narrow down the number of candidate point mutations from ∼35 million differences since the split from chimpanzee when comparing only reference genomes (Consortium 2005) to 31,389 fixed human-specific changes in a previous seminal study (Pääbo 2014). Other types of more complex changes like structural variants most likely contributed to human-specific traits. For example, it is well known that since the split from chimpanzees functional differences arose through gene duplications in *ARHGAP11B* and other genes (Florio et al. 2015; Ju et al. 2016), copy number variants in *SRGAP2* and other genes (Dennis et al. 2012; Dumas et al. 2012; Suzuki et al. 2018) or regulatory deletions (McLean et al. 2011). In these cases, the variants arose before the split of humans and Neanderthals, but the differences in structural variation that exist between the hominin lineages (Chintalapati et al. 2017) need to be explored in more detail, with advancement of technologies in ancient DNA sequencing and computational methods. This will result in complementary lists of changes for understanding the human condition outside the scope of this study. Beyond that, parts of the genome which are complex and not yet examined by conventional sequencing platforms (O’Bleness et al. 2012) possibly harbor important human-specific changes as well.

**Figure 1:**
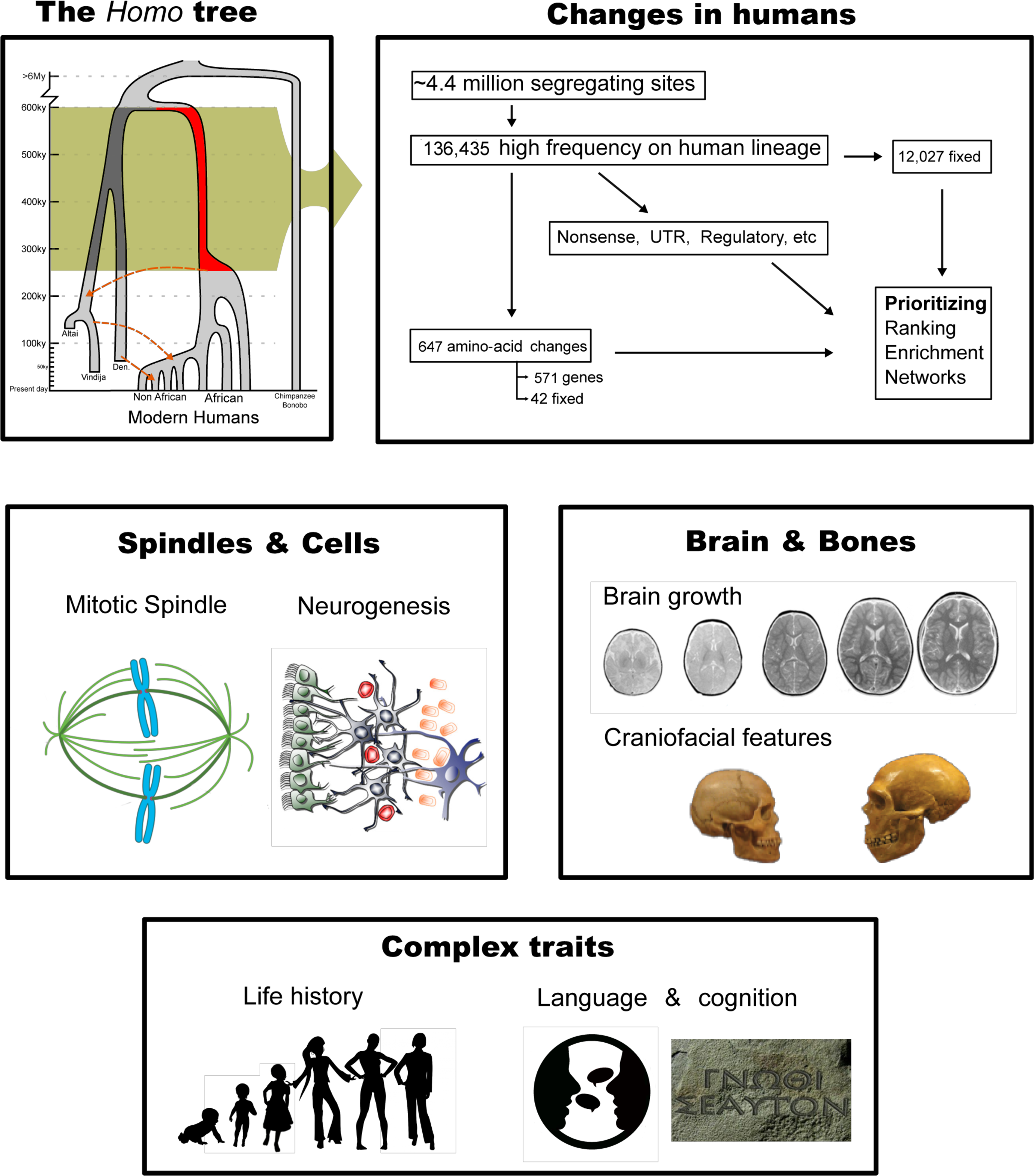
Conceptual summary of this study.

Some of the single nucleotide changes have been linked to putative functional consequences (Castellano et al. 2014; Pääbo 2014; Prüfer et al. 2014), and evidence is mounting that several molecular changes affecting gene expression in the brain were subject to selective pressures (Green et al. 2010; Somel et al. 2013; Zhou et al. 2015; Racimo 2016; Peyrégne et al. 2017). Furthermore, the genomic impact of interbreeding events is not evenly distributed across the genome. Genes expressed in regions of the brain regarded as critical for certain cognitive functions such as language are depleted in introgressed archaic genetic material (Sankararaman et al. 2014; Vernot and Akey 2014; Sankararaman et al. 2016; Vernot et al. 2016), and introgressed alleles are downregulated in some of these brain regions, suggesting natural selection acting on tissue-specific gene regulation (McCoy et al. 2017). Thus, it seems reasonable to conclude that there were differences between anatomically modern human and Neanderthal brains, and that these underlie at least some of the characteristics of our lineage (Wynn and Coolidge 2004). We want to emphasize that such recent differences are likely to be subtle when compared to those after the split from our closest living relatives on a scale of 6-10 million years (Langergraber et al. 2012), where fundamental changes arose since the divergence from chimpanzees and bonobos (Varki and Altheide 2005; O’Bleness et al. 2012). The observation of recurrent gene flow between modern human and archaic populations also implies a broad overall similarity, yet, such subtle differences may still have contributed to the evolutionary outcome (Wynn et al. 2016). This does not imply a superiority of humans, but specific changes that might have facilitated survival under the given environmental conditions. Obviously, not all human-specific changes are beneficial: While most mutations may be rather neutral and have little effect on the phenotype, some may have had deleterious effects or side-effects, possibly increasing the risks for neurodevelopmental or neurodegenerative disorders in humans (Bufill et al. 2011; Bruner and Jacobs 2013; Bufill et al. 2013).

The goal of this paper is to provide a revised, extended set of recent single nucleotide changes in humans since their split from Neanderthals that could enrich our understanding of the molecular basis of recent human condition. The previous focus on fixed alleles was reasonable given limited data (Pääbo 2014), but having a better grasp of the magnitude of modern human variation and the interaction between different hominin lineages seems a good reason to cast a wider net, and take into account not only fixed differences but also high-frequency changes shared by more than 90% of present-day individuals. Here, we present a revised list of 36 genes that carry missense substitutions which are fixed across thousands of human individuals and for which all archaic hominin individuals sequenced so far carry the ancestral state. In total, 647 protein-altering changes in 571 genes reached a frequency of at least 90% in the present-day human population. We attempt to interpret this list, as well as some regulatory changes, since it seems very likely that some of these genes would have contributed to the human condition. We discuss some of their known functions, and how these relate to pathways that might have been modified during human evolution from the molecular level to cellular features and more complex phenotypic traits (Fig. 1). We restrict our attention to genes where the literature may allow firm conclusions and predictions about functional effects, since many genes likely have multiple different functions (Gratten and Visscher 2016). Obviously, it cannot be emphasized enough that ultimately, experimental validation will be needed to confirm our hypotheses concerning alterations in specific functions. For example, transcription factors or enzymatically active proteins can be tested using cell cultures or *in vitro* assays, while brain organoids could be used to test differences in neuronal functions (Giandomenico and Lancaster 2017), especially in combination with single-cell RNA sequencing (Camp et al. 2015; Camp and Treutlein 2017). Ultimately, these variants can be introduced into model organisms like mice to test complex features related to cognitive abilities or behavior (Enard et al. 2009). Still, given limitations to the amount of changes that can be tested at once, networks which are modified by multiple changes cannot be tested with current technologies.

## Results

### Genetic differences between present-day humans and archaic hominins

Using publicly available data on one Denivosan and two Neanderthal individuals and present-day human variation (Methods), we calculated the numbers of single nucleotide changes (SNCs) which most likely arose recently on the respective lineages after their split from each other, and their functional consequences (Table 1). Previously, a number of 31,389 sites has been reported as recently fixed derived in present-day humans, while being ancestral in archaics (Pääbo 2014; Prüfer et al. 2014). We find a smaller number of only 12,027 positions in the genome, in part because of including another archaic individual and different filters, but mainly by a richer picture of present-day human variation. The 1,000 Genomes Project as well as other sources contributing to the dbSNP database now provide data for thousands of individuals, which results in very high allele frequencies for many loci instead of fixation. Indeed, 29,358 positions show allele frequencies larger than 0.995, demonstrating that the level of near-fixation is similar to the level of previously presented fixation. The number of loci with high frequency (HF) changes of more than 90% in present-day humans is an order of magnitude larger than the number of fixed differences. The three archaic individuals carry more than twice as many changes than present-day humans; however, we emphasize that much of this difference is not due to more mutations in archaics, but rather the fact that data for only three individuals is available, compared to thousands of humans. The variation across the archaic population is not represented equally well, which makes these numbers not directly comparable. On the other hand, much less variation is found by the sequencing of each additional Neanderthal individual compared to humans due to the low diversity of Neanderthals (Fig. S36 in (Kuhlwilm et al. 2016)). This low diversity across their geographic range suggests that most alleles observed as ancestral here will be the same state in other individuals. Furthermore, we take variability into account due to gene flow or errors, decreasing the possibility that positions ancestral in the archaic individuals studied to date turn out to be derived in most archaic individuals, hence this extended catalog will likely not undergo drastic changes. However, changes in structural variants or regions of the genome that are not accessible by current sequencing technologies will most likely complement our results (O’Bleness et al. 2012).

**Table 1:** Summary of single nucleotide changes. TFBS: Transcription factor binding sites. UTR: Untranslated Region. HF: High frequency. Fixed changes are a subset of HF changes.

|  | **Fixed human** | **HF human** | **Extended human** | **Fixed archaic** | **HF archaic** | **Extended archaic** |
| --- | --- | --- | --- | --- | --- | --- |
| **All** | 12,027 | 136,435 | 83,254 | 33,498 | 380,756 | 983 |
| **Non-synonymous** | 42 | 647 | 327 | 167 | 1,921 | 13 |
| **Synonymous** | 41 | 843 | 363 | 193 | 2,123 | 14 |
| **Start/stop** | 1 | 14 | 10 | 3 | 48 | 2 |
| **Splice site** | 4 | 23 | 8 | 4 | 54 | 0 |
| **TFBS** | 28 | 226 | 126 | 87 | 914 | 1 |
| **Upstream** | 1,935 | 19,599 | 11,235 | 4,920 | 55,188 | 289 |
| **5' UTR** | 180 | 1,853 | 1,012 | 195 | 2,016 | 7 |
| **3' UTR** | 77 | 702 | 334 | 506 | 5,303 | 19 |
| **Downstream** | 1,922 | 19,704 | 11,673 | 4,956 | 55,832 | 281 |
| **miRNA** | 0 | 1 | 2 | 0 | 4 | 0 |
| **Regulatory element** | 1,952 | 20,971 | 12,320 | 5,125 | 59,248 | 195 |

Present-day humans carry 42 fixed amino acid-changes in 36 genes (Table 2, Fig. 2), while Neanderthals carry 159 such changes. Additionally, modern humans carry 605 amino acid-changes at high frequency (human-lineage high-frequency missense changes, referred to as HHMCs), amounting to a total of 647 such changes in 571 genes (Table S1). Together with 323 SNCs on the human lineage with low confidence (Methods, Table S2), almost 1,000 putative protein-altering changes were found across most present-day humans. Generally, synonymous changes are found at a similar magnitude as missense changes, but only few SNCs altering start and stop codons, and thousands of changes in putative regulatory and untranslated regions. We admit that some of the loci presented here are variable across the phylogenetic tree, or less reliable due to low coverage in the archaics, but we accept this since our intention is retrieve an inclusive picture of possibly functional recent changes. The 42 protein-altering changes for which the ancestral allele has not been observed in any present-day human, most of which have been presented before (Pääbo 2014), constitute without doubt the strongest entry points into a molecular understanding of the human condition, and should be prime candidates for experimental validation. Only one gene, *SPAG5,* carries three such SNCs, and four genes (*ADAM18, CASC5, SSH2* and *ZNHIT2*) carry two fixed protein-coding changes in all modern humans. We identified 15 SNCs (in *AHR, BOD1L1, C1orf159, C3, DNHD1, DNMT3L, FRMD8, OTUD5, PROM2, SHROOM4, SIX5, SSH2, TBC1D3, ZNF106, ZNHIT2*) that have not been previously described as fixed differences between humans and archaics. We note that another 12 previously described (Pääbo 2014) protein-altering substitutions were not found among the genotypes analyzed here (in *C21orf62, DHX29, FAM149B1, FRRS1L, GPT, GSR, HERC5, IFI44L, KLF14, PLAC1L, PTCD2, SCAF11*). These genotype calls are absent from the files provided for the three archaic genomes due to different genotype calling and filtering procedures compared to the original publication of the Altai Neanderthal genome (Prüfer et al. 2014; Prüfer et al. 2017). Hence, some potentially relevant candidate changes were not included here, and future research is necessary to evaluate these as well. Despite attempting an extended interpretation, our data is thus not fully exhaustive.

**Table 2:**
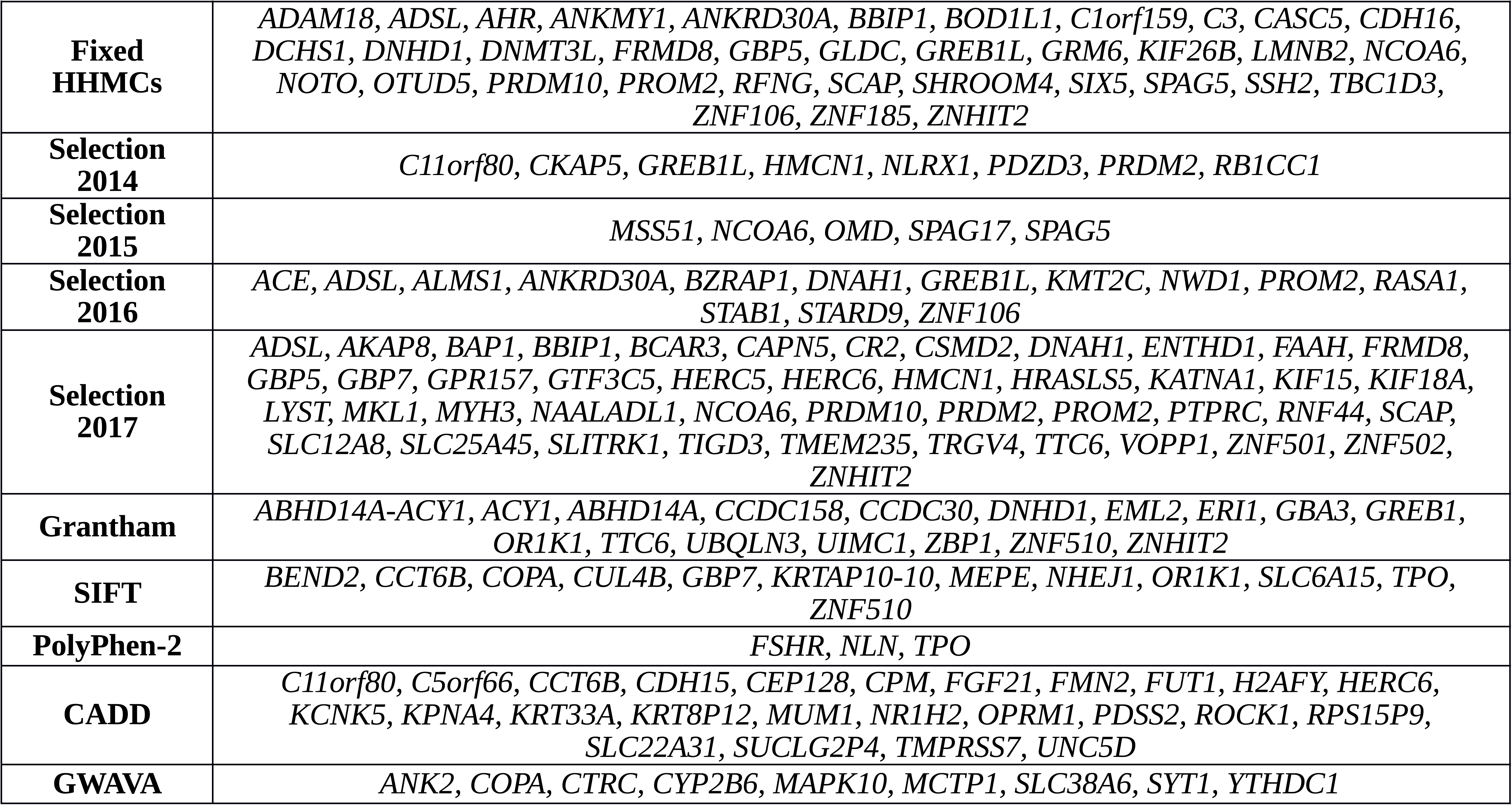
Genes with fixed non-synonymous changes on the human lineage, genes under positive selection with HHMCs, and deleterious candidate HHMCs. Selection 2014: Prüfer *et al.*, 2014. Selection 2015: Zhou *et al.*, 2015. Selection 2016: Racimo, 2016. Selection 2017: Peyrégne *et al.*, 2017.

**Figure 2:**
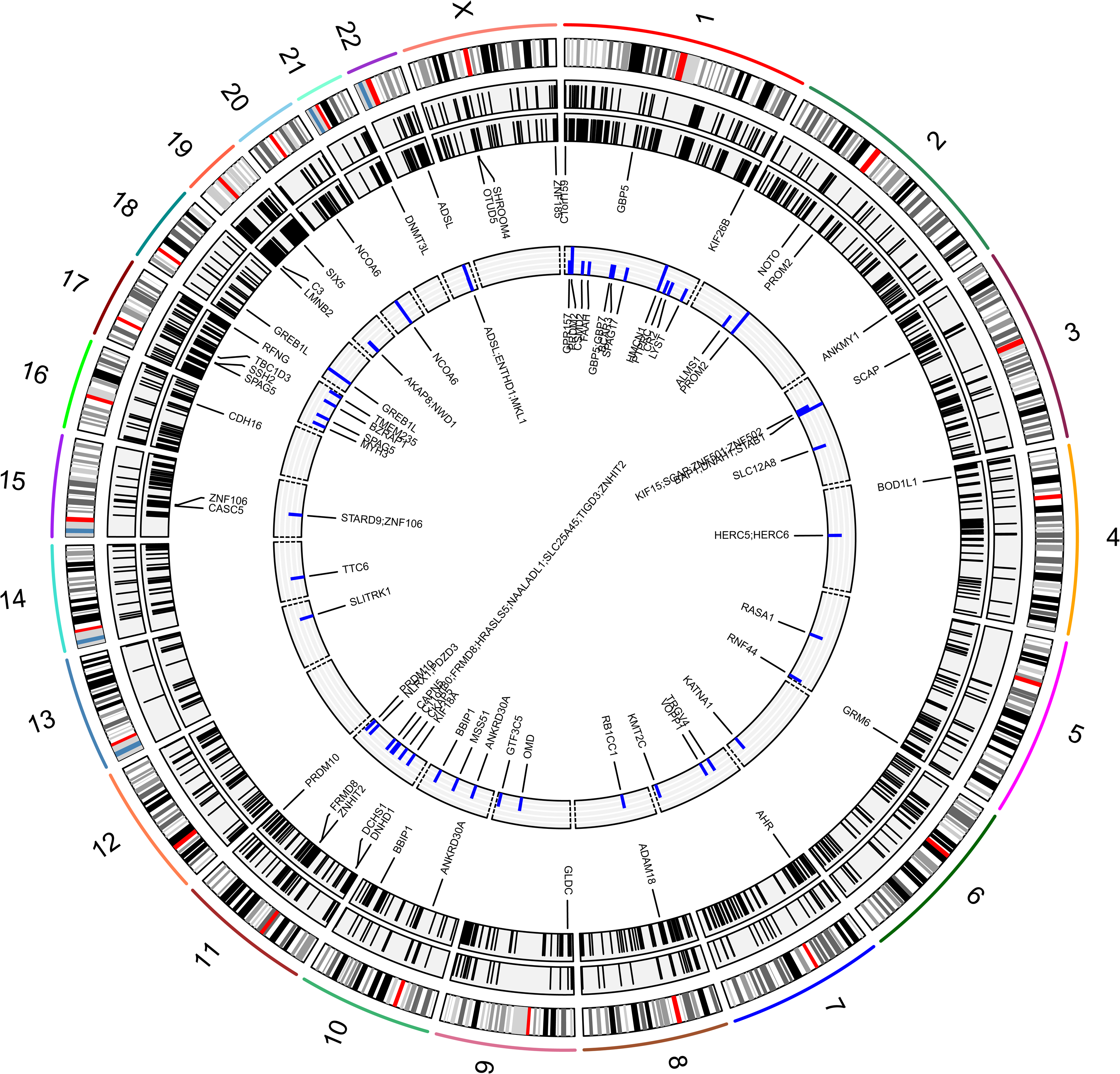
Features discussed in this study. From inside to outside: Genes with HHMCs and signatures of positive selection (compare Table 2), genes with fixed non-synonymous SNCs on the human lineage, HHMCs, AHMCs, karyogram of human chromosomes.

It is noteworthy that the number of fixed SNCs decreased substantially, and it is possible that single individuals will be found to carry some of the ancestral alleles for the remaining fixed sites. Hence, it is important to focus not only on fixed differences, but also consider variants at high frequency. When analyzing the 647 HHMCs, 68 genes carry more than one amino acid-altering change. Among these, *TSGA10IP* (Testis Specific 10 Interacting Protein) and *ABCC12* (ATP Binding Cassette Subfamily C Member 12) carry four such changes, and seven more genes (*MUC5B, NPAP1, OR10AG1, OR5M9, PIGZ, SLX4, VCAN*) carry three HHMCs. 1,542 genes carry at least one HF missense change on the archaic lineage (archaic-lineage high-frequency missense change, referred to as AHMC, Tables S3, S4). We find an overlap of 122 genes with HHMCs and AHMCs, which is more than expected considering that among 1,000 sets of random genes of a similar length distribution, no overlap of this extent was observed. The same genes seem to have acquired missense changes on both lineages since their divergence more often than expected. We find a high ratio of HHMCs over synonymous changes for chromosome 21 (1.75-fold), and a very small ratio (0.18-fold) for chromosome 13. We do not find such extreme ratios for AHMCs and corresponding synonymous changes, suggesting differences in the distribution of amino acid changes between both lineages (Fig. S1).

### Ranking and enrichment

We assessed the impact of mutations for different deleteriousness scores (Table 2), finding 12 genes with deleterious HHMCs according to SIFT, three according to PolyPhen, and 16 when using the Grantham score (>180), measuring the physical properties of amino acid changes. The C-score and GWAVA can be used to rank all mutation classes, and we present the top candidates.

Then, we attempted a ranking of genes by the density of lineage-specific changes in the dataset. As expected, the total number of segregating sites is correlated with gene length (Pearsons’ R = 0.93). This correlation is weaker for HF human SNCs (R = 0.73) and fixed human-specific SNCs (R= 0.25), as well as for fixed (R = 0.37) and HF (R = 0.82) SNCs in archaics. We conclude that some genes with a large number of human-specific changes might carry these large numbers by chance, while others are depleted. Indeed, 17,453 (88.9%) of these genes do not carry any fixed human-specific change, and 80.5% do not carry fixed archaic-specific changes. Of note, genes that have attracted attention in the context of traits related to the “human condition” like *CNTNAP2* and *AUTS2* are among the longest genes in the genome, hence changes in these genes should be interpreted with caution as they are not unexpected. We ranked the genes by the number of HF changes in either modern humans or archaics, divided by their genomic lengths, and categorize the top 5% of this distribution as putatively enriched for changes on each lineage (Table S5). We note that 191 genes (30.9%) fall within this category for both human HF changes and archaic HF changes, as a result of differences in mutation density. In order to distinguish a truly lineage-specific enrichment, we calculated the ratios of HF changes for humans and archaics, defining the top 10% of genes in this distribution as putatively enriched (Table S5). Among the genes enriched for changes on the modern human lineage, 18 carry no HF changes on the archaic lineage, and ten of these also fall within the 5% of genes carrying many changes considering their length (*ARSJ, CLUAP1, COL20A1, EPPIN, KLHL31, MKNK1, PALMD, RIC3, TDRD7, UBE2H*). These might be candidates for an accumulation of changes, even though this is not identical to selective sweep signals. Among these, the collagen COL20A1 and the Epididymal Peptidase Inhibitor EPPIN carry HHMCs. *ACAD10, DST* and *TTC40,* which carry two HHMCs, might be other notable genes with a human-specific enrichment.

Gene Ontology (GO) categories are neither enriched for HHMCs on the human lineage in a hypergeometric test, nor for genes carrying AHMCs, HF changes in UTRs or transcription factor binding sites. However, instead of singular changes that might be observed more often in long genes, or genes that are more prone to mutations in hominins, the density of HF changes in a gene might yield a better picture of lineage-specific changes, possibly for cumulative changes. We applied a test for the ratio of the number of gene-wise HF changes on one lineage over the other lineage, finding an enrichment for 12 GO categories on the human lineage (Table S6), with “soft palate development”, “negative regulation of adenylate cyclase activity”, “collagen catabolic process” and “cell adhesion” in the biological process category. Among the cellular components category, the “postsynaptic membrane”, “spermatoproteasome complex”, “collagen trimer”, “dendrite” and “cell junction” show enrichment, as well as the molecular functions “calcium ion binding”, “histone methyltransferase activity (H3-K27 specific)” and “metallopeptidase activity”. We find no GO enrichment for genes with an excess of changes on the archaic lineage. In order to approach a deeper exploration of genes with associated complex traits in humans, we explored the NHGRI-EBI GWAS Catalog (MacArthur et al. 2017), containing 2,385 traits. We performed a systematic enrichment screen, finding 17 unique traits enriched for genes with HHMCs, and 11 for genes with AHMCs (Table S7). Changes in genes associated to “Cognitive decline (age-related)”, “Rheumatoid arthritis” or “Major depressive disorder” might point to pathways that could have been influenced by protein-coding changes on the human lineage. In archaics, genes are enriched, among others, for associations to traits related to body mass index or cholesterol levels, which might reflect differences in their physiology.

We find a significant enrichment of protein-protein interactions (P = 0.006) among the gene products of HHMC genes (Fig. S2), meaning that these proteins interact with each other more than expected. Functional enrichment is found for the biological process “cellular component assembly involved in morphogenesis”, most strongly for the cellular components cytoskeleton and microtubule, as well as the molecular function “cytoskeletal protein binding”. Three proteins have at least 20 interactions in this network and might be considered important nodes: TOP2A, PRDM10 and AVPR2 (Table S8). However, proteins encoded by genes with synonymous changes on the modern human lineage seem to be enriched for interactions as well (P = 0.003), as are proteins encoded by genes with AHMCs (P = 1.68 x 10^-14^), with an enrichment in GO categories related to the extracellular matrix and the cytoskeleton, and proteins with more than 40 interactions (Table S8). We caution that these networks might be biased due to more mutations and possibly more interactions in longer, multi-domain genes.

Regulatory changes might have been important during our evolution (Wray 2007), hence we tested for an overrepresentation of transcription factors. We find 78 known or putative transcription factors among the HHMC genes (Table S9) on the modern human lineage (Chawla et al. 2013), which is not overrepresented among genes with HHMCs (with 49.2% of random genes sets containing fewer HHMCs). Despite no enrichment as a category, single transcription factors on the modern human lineage might have been important, particularly those with an excess of modern human over archaic HF changes (*AHR, MACC1, PRDM2, TCF3, ZNF420, ZNF516*). Others, like *RB1CC1* (Prüfer et al. 2014) or *PRDM10* and *NCOA6* (Peyrégne et al. 2017) have been found in selective sweep screens, suggesting contributions of individual transcription factors, rather than the class of proteins. We also tested for an enrichment of gene expression in different brain regions and developmental stages (Miller et al. 2014; Grote et al. 2016), using the HF synonymous changes on each lineage as background sets. We find an enrichment of gene expression in the orbital frontal cortex at infant age (0-2 years) for genes with HHMCs, but no enrichment for genes with AHMCs. Furthermore, when testing the genes with HHMCs and using the set of genes with AHMCs as background, “gray matter of forebrain” at adolescent age (12-19 years) is enriched, while no enrichment was found for genes with AHMCs.

## Discussion

The enrichment of broad categories above suggests that traits related to brain functions are prominently represented by HHMCs. It should be noted that such results would be less clear if we just focused on completely fixed changes, given the drastically reduced number of genes harboring such changes. Here, we will further examine the possible impact on the brain that some of these changes might have, paying special attention to hypotheses formulated in earlier work on modern human-specific changes. Our extended catalog of changes appears to provide additional support for some of these hypotheses.

### Cell division and the brain growth trajectory

It has been proposed previously that protein-coding changes in cell cycle-related genes are highly relevant candidates for human-specific traits (Pääbo 2014; Prüfer et al. 2014). Indeed, three genes (*CASC5, SPAG5*, and *KIF18A*) have been singled out as involved in spindle pole assembly during mitosis (Pääbo 2014). Other genes with protein-coding SNCs (*NEK6* and *STARD9/KIF16A*) turn out to be implicated in the regulation of spindle pole assembly as well (O’Regan and Fry 2009; Torres et al. 2017). Among the 15 fixed protein-coding changes identified here but absent from previous analyses (Pääbo 2014; Prüfer et al. 2014), some might have also contributed to complex modifications of pathways in cell division, like *AHR* (Puga et al. 2002) or *DNHD1* (Bader et al. 2011) (Supplementary Information 1), as well as other genes with HHMCs, like *CHEK1* (Zachos et al. 2017) or the gene encoding for the protein TOP2A (Yoshida and Azuma 2016), which shows the largest number of interactions with other HHMC-carrying proteins, suggesting a function as interaction hub in the cell division complex (Supplementary Information 1). Taken together, these changes suggest that the cell cycle machinery might have been modified in a specific way in humans compared to other hominins.

It has been claimed (Prüfer et al. 2014) that genes with fixed non-synonymous changes in humans are also more often expressed in the ventricular zone of the developing neocortex, compared to fixed synonymous changes. Since the kinetochore-associated genes *CASC5, KIF18A* and *SPAG5* are among these genes, it has been emphasized that this “may be relevant phenotypically as the orientation of the mitotic cleavage plane in neural precursor cells during cortex development is thought to influence the fate of the daughter cells and the number of neurons generated (Fietz and Huttner 2011)” (Prüfer et al. 2014). Several fixed SNCs on the modern human lineage are observed for *CASC5* (two changes) and *SPAG5* (three changes), which is also among genes with a relatively high proportion of HF changes (Table S5). The changes in *KIF18A, KIF16A* and *NEK6* can no longer be considered as fixed, but occur at very high frequencies (>99.9%) in present-day humans. We attempted to determine whether an enrichment of genes with HHMCs on the human lineage can be observed in the ventricular zone (Miller et al. 2014), but instead find an enrichment in the intermediate zone, where less than 5% of random gene sets of the same size are expressed. However, synonymous HF changes also show an enrichment in this layer, as well as genes with AHMCs (Table S10), suggesting an overrepresentation of genes that carry mutations in the coding regions rather than lineage-specific effects. We were able to broadly recapitulate the observation of an enrichment of expression in the ventricular zone if restricting the test to genes with non-synonymous changes at a frequency greater than 99.9% in present-day humans, which is not observed for corresponding synonymous and archaic non-synonymous changes (Table S10). Among the 28 genes expressed in the ventricular zone that carry almost fixed HHMCs, four might be enriched for HF changes in humans (*HERC5, LMNB2, SPAG5, VCAM1*), and one shows an excess of HF changes on the human compared to the archaic lineage (*AMKMY1*). Other notable genes discussed in this study include *ADSL, FAM178A, KIF26B, SLC38A10*, and *SPAG17*.

The centrosome-cilium interface is known to be critical for early brain development, and centrosome-related proteins are overrepresented in studies on the microcephaly phenotype in humans (Megraw et al. 2011). We find 126 genes (Table S9) with 143 HHMCs that putatively interact with proteins at the centrosome-cilium interface (Gupta et al. 2015). Some of the genes listed here and discussed in this study, such as *FMR1, KIF15, LMNB2, NCOA6, RB1CC1, SPAG5* and *TEX2,* harbor not only HHMCs, but an overall high proportion of HF changes on the human lineage. Although an early analysis suggested several candidate genes associated to microcephaly, not all of these could be confirmed by high-coverage data. Among eleven candidate genes (Green et al. 2010), only two (*PCNT, UCP1*) are among the HHMC gene list presented here, while most of the other changes are not human-specific, and only *PCNT* has been related to microcephaly (Li et al. 2015). Nevertheless, more changes related to microcephaly are found on both lineages, for example in *ATRX* (K. Ritchie et al. 2014) or *CASC5* (Genin et al. 2012) (Supplementary Information 3).

Changes in genes associated with brain growth trajectory differences lead not necessarily to a decrease but also an increase of brain size (Montgomery et al. 2011), suggesting that the disease phenotype of macrocephaly might point to genes relevant in the context of brain growth as well. One of the few genes with several HHMCs, *CASC5*, has been found to be associated with gray matter volume differences (Shi et al. 2017). It has been claimed that mutations in *PTEN* alter the brain growth trajectory and allocation of cell types through elevated Beta-Catenin signaling (Chen et al. 2015). This well-known gene, critical for brain development (Li et al. 2017), has not been highlighted in the context of human-specific changes, while we find that *PTEN* falls among the genes with an excess on the modern human over the archaic lineage, suggesting that regulatory changes in this gene might have contributed to human-specific traits. This is also the case for the HHMC-carrying transcription factor TCF3, which is known to repress Wnt-Beta-Catenin signaling and maintain the neural stem cell population during neocortical development (Kuwahara et al. 2014). Changes in these and other genes (Supplementary Information 3) like *CCND2* (Mirzaa et al. 2014), *GLI3* (Jamsheer et al. 2012), or *RB1CC1* (Wang et al. 2013), for which a regulatory SNC has been suggested to modify transcriptional activity (Weyer and Pääbo 2016) and which carries a signature of positive selection (Prüfer et al. 2014), could have contributed to the brain growth trajectory changes hypothesized to give rise to the modern human-specific globular braincase shape during the past several 100,000 years (Gunz et al. 2012; Hublin et al. 2015; Neubauer et al. 2018). Finally, we find changes that might have affected the size of the cerebellum, a key contributor to our brain shape (Kochiyama et al. 2018; Neubauer et al. 2018), such as HF regulatory SNCs in *ZIC1* and *ZIC4* (Blank et al. 2011), an excess of HF mutations in *AHI1* (Cheng et al. 2012), and a deleterious HHMC in *ABHD14A*, which is a target of ZIC1 (Hoshino et al. 2003).

### Cellular features of neurons

To form critical networks during the early development of the brain, axonal extensions of the neurons in the cortical region must be sent and guided to eventually reach their synaptic targets. Studies conducted on avian vocal learners (Pfenning et al. 2014; Wang et al. 2015) have shown a convergent differential regulation of axon guidance genes of the *SLIT-ROBO* families in the pallial motor nucleus of the learning species, allowing for the formation of connections virtually absent in the brains of vocal non-learners. In modern humans, genes with axon-guidance-related functions such as *FOXP2, SLIT2* and *ROBO2* have been found to lie within deserts of archaic introgression (Sankararaman et al. 2016; Vernot et al. 2016; Kuhlwilm 2018), suggesting incompatibilities between modern humans and archaics for these regions. Our dataset contains a fair amount of genes known to impact brain wiring: Some of the aforementioned microtubule-related genes, specifically those associated with axonal transport and known to play a role in post-mitotic neural wiring and plasticity (Lüders 2016), are associated with signals of positive selection, such as *KIF18A* (McVicker et al. 2016) or *KATNA1* (Ahmad et al. 1999; Karabay et al. 2004). Furthermore, an interactor of KIF18A, KIF15 (Kevenaar et al. 2017), might have been under positive selection in modern humans (Peyrégne et al. 2017), and contains two HHMCs. Versican (*VCAN*), which promotes neurite outgrowth (Wu et al. 2004), carries three HHMCs, and *SSH2* (two HHMCs) might be involved in neurite outgrowth (Cuberos et al. 2015). *PIEZO1*, which carries a non-synonymous change that is almost fixed in modern humans, is another factor in axon guidance (Koser et al. 2016), as well as *NOVA1* (Jensen et al. 2000), which is an interactor of *ELAVL4* (Ratti et al. 2008), a gene that codes for a neuronal-specific RNA-binding protein and might have been under positive selection in humans (Zhou et al. 2015; Peyrégne et al. 2017). Furthermore, we find one of the most deleterious regulatory SNCs in the Netrin receptor UNC5D, which is critical for axon guidance (Takemoto et al. 2011).

We also detect changes in genes associated with myelination and synaptic vesicle endocytosis, critical to sustain a high rate of synaptic transmission, including *DCX* (Yap et al. 2012), *SCAP* (Verheijen et al. 2009), *RB1CC1* (Menzies et al. 2015), *ADSL* (Jurecka et al. 2012) and *PACSIN1* (Widagdo et al. 2016) among others (Supplementary Information 2). It is noteworthy that among traits associated with cognitive functions such as language or theory of mind, the timing of myelination appears to be a good predictor of computational abilities (Skeide and Friederici 2016; Grosse Wiesmann et al. 2017). Computational processing might have been facilitated by some of the changes presented here, at least in some of the circuits that have expanded in our lineage (Mars et al. 2018), since subtle maturational differences early in development (Dubois et al. 2016) may have had a considerable impact on the phenotype. In this context, it is worth mentioning that in our dataset, several genes carrying HHMCs and associated with basal ganglia functions (critical for language and cognition) stand out, like *SLITRK1* (Abelson et al. 2005) and *NOVA1* (Jelen et al. 2010; Konopka et al. 2012; Alsiö et al. 2013; Zhou et al. 2015; Popovitchenko et al. 2016) (Supplementary Information 4). Finally, in the broader context of cognition, we find an enrichment of HHMCs in genes associated to “Alzheimer’s disease (cognitive decline)” and “Cognitive decline (age-related)”, with seven associated genes (*COX7B2, BCAS3, DMXL1, LIPC, PLEKHG1, TTLL2* and *VIT*). Among genes influencing behavioral traits (Supplementary Information 4) are *GPR153* (Sreedharan et al. 2011), *NCOA6* (Takata et al. 2018), or the Adenylosuccinate Lyase (*ADSL*) (Fon et al. 1995), for which the ancestral Neanderthal-like allele has not been observed in 1,000s of modern human genomes and which has been pointed out before as under positive selection (Castellano et al. 2014; Racimo et al. 2014; Racimo 2016; Peyrégne et al. 2017) We know that archaic hominins likely had certain language-like abilities (Dediu and Levinson 2013; Dediu and Levinson 2018), and hybrids of modern and archaic humans must have survived in their communities (Fu et al. 2015), underlining the large overall similarity of these populations. However, genes associated with axon guidance functions, which are important for the refinement of neural circuits including those relevant for speech and language, are found in introgression deserts (Jeong et al. 2016; Lei et al. 2017), which seems to be a unidirectional and human-specific pattern especially in the *FOXP2* region (Kuhlwilm 2018). We suggest that modifications of a complex network in cognition or learning took place in modern human evolution (Boeckx and Benítez-Burraco 2014), possibly related to other brain-related (Bastir et al. 2011; Hublin et al. 2015; Boeckx 2017; Bryant and Preuss 2018), vocal tract (Gokhman et al. 2017) or neural changes (Belyk and Brown 2017).

### The craniofacial phenotype

In previous work on ancient genomes changes related to craniofacial morphology have been highlighted (Castellano et al. 2014; Gokhman et al. 2017), and we find an enrichment of genes with an excess of HF SNCs on the modern human lineage for soft palate development (Table S6). Among genes harboring an excess of HF SNCs associated with specific facial features, we find *RUNX2, EDAR,* and *GLI3* (Adhikari et al. 2016), *NFATC1* (Kim and Kim 2014), *SPOP* (Cai and Liu 2016), *DDR2* (Zhang et al. 2011) and *NELL1* (Zhang et al. 2012), possibly carrying changes in regulatory regions, while mutations in the HHMC-carrying gene encoding for the transcription factor ATRX cause facial dysmorphism (Moncini et al. 2013). In addition, genes with HHMCs such as *PLXNA2* (Oh et al. 2012), *EVC2* (Kwon et al. 2018), *MEPE* (Gullard et al. 2016), *OMD* (Tashima et al. 2015), and *SPAG17* (Teves et al. 2015) are known to affect craniofacial bone and tooth morphologies. These genes appear to be important in determining bone density, mineralization and remodeling, hence they may underlie differences between archaic and modern human facial growth (Lacruz et al. 2015). Some of these facial properties may have been present in the earliest fossils attributed to *H. sapiens,* like the Jebel Irhoud fossils (Hublin et al. 2017), deviating from craniofacial features which emerged in earlier forms of *Homo* (Lacruz et al. 2013), and may have become established before some brain-related changes discussed here (Stringer 2016; Neubauer et al. 2018). The gene encoding the transcription factor PRDM10 stands out for carrying HHMCs, being found in selective sweep regions and the second-most interacting protein within the HHMC dataset. Although little is known about *PRDM10*, it may be related to dendrite growth (Siegel et al. 2002) and neural crest related changes that contributed to the formation of our distinct modern face (Park and Kim 2010). Other craniofacial morphology-related genes, such as *DCHS2* (Adhikari et al. 2016), *HIVEP2* (Jones et al. 2010), *HIVEP3* (Imamura et al. 2014), *FREM1* (Lee et al. 2017), and *FRAS1* (Talbot et al. 2016) harbor AHMCs, while another bone-related gene, *MEF2C* (Verzi et al. 2007), shows an excess of HF changes on the archaic lineage. These changes may underlie some of the prominent derived facial traits of Neanderthals (Rak 1986).

### Life history and other phenotypic traits

Apart from their consequences for cognitive functions, it has been suggested that changes involved in synaptic plasticity might be interpreted in a context of neoteny (Somel et al. 2009; Liu et al. 2012; Peyrégne et al. 2017; Sherwood and Gómez-Robles 2017), with the implication of delayed maturation in humans (Bednarik 2013) and a longer timeframe for brain development. However, given their similar brain sizes (Hofman 1983), humans and Neanderthals might both have needed a long overall maturation time (Ponce de León et al. 2017; Rosas et al. 2017). Accordingly, notions like neoteny and heterochrony are unlikely to be fine-grained enough to capture differences between these populations, but early differences in infant brain growth between humans and Neanderthals (Gunz et al. 2010; Hublin et al. 2015) could have rendered our maturational profile distinct during limited developmental periods and within specific brain regions, imposing different metabolic requirements (Bruner et al. 2014). One of the brain regions where such differences are found is the orbitofrontal cortex (OFC) (Bastir et al. 2011), and we find that the OFC at infant age (0-2 years) is enriched for the expression of genes that carry HHMCs compared to synonymous SNCs. We suggest that the development of the OFC in infants might have been subject to subtle changes since the split from Neanderthals rather than a general developmental delay, which is particularly interesting given that this brain region has been implied in social cognition (Beer et al. 2006) and learning (Miller et al. 2018).

Genes carrying HHMCs are enriched for expression in the gray matter of the forebrain at the adolescent age compared to AHMC-carrying genes, hence additional human-specific modifications during this period might have taken place, possibly linked to changes in myelination described above. It has been suggested that differences in childhood adolescence time existed between humans and Neanderthals, after a general developmental delay in the hominin lineage (Smith and Tompkins 1995; Bock and Sellen 2002). Dental evidence suggests an earlier maturation in Neanderthals than modern humans (Smith et al. 2010), and it has been claimed that Neanderthals might have reached adulthood earlier (Ramirez Rozzi and de Castro 2004). Furthermore, an introgressed indel from Neanderthals causes an earlier onset of menarche in present-day humans (Chintalapati et al. 2017), supporting at least the existence of alleles for earlier maturation in the Neanderthal population. Among the genes carrying fixed HHMCs, *NCOA6* has also been linked to age at menarche and onset of puberty (Day et al. 2017), as well as placental function (Antonson et al. 2003). This putative transcription factor is enriched in HF changes and has been suggested to have been under positive selection on the modern human lineage (Racimo et al. 2014; Peyrégne et al. 2017). The HHMC is located nearby and three 5’-UTR variants within a putatively selected region (Zhou et al. 2015), with an estimated time of selection at around 150 kya (assuming a slow mutation rate). Even though this gene carries an AHMC as well, it remains possible that modern humans acquired subtle differences in their reproductive system through lineage-specific changes in this gene. A delay in reproductive age may influence overall longevity, another trait for which our data set yields an enrichment of genes with HHMCs (*SLC38A10, TBC1D22A* and *ZNF516*).

The male reproductive system might have been subject to changes as well, since we find that several proteins in spermatogenesis seem to carry two HHMCs: Sperm Specific Antigen 2 (*SSFA2*), Sperm Associated Antigen 17 (*SPAG17*), *ADAM18* (Zhu et al. 1999) and *WDR52* (Tang et al. 2017), out of which *ADAM18* and *SPAG17* also carry AHMCs. Lineage-specific differences in genes related to sperm function or spermatogenesis might have been relevant for the genetic compatibility between humans and Neanderthals. Another gene harboring a HHMC with similar functions is *EPPIN* (Wang et al. 2005), which shows no HF changes on the archaic, but 27 such SNCs on the modern human lineage. The gene encoding for the Testis Expressed 2 protein (*TEX2*) is enriched for HF changes in both humans and archaics, with one HHMC and five AHMCs, but its function is not yet known. Another possible SNC that might be relevant in this context is a splice site change in *IZUMO4*, since proteins encoded by the IZUMO family form complexes on mammalian sperm (Ellerman et al. 2009). The adjacent exon is not present in all transcripts of this gene, suggesting a functional role of this splice site SNC. Finally, genes in the GO category “spermatoproteasome complex” are enriched for an excess of HF changes on the human lineage.

It has been found that Neanderthal alleles contribute to addiction and, possibly, pain sensitivity in modern humans (Simonti et al. 2016; Dannemann et al. 2017). In this context, an interesting protein-truncating SNC at high frequency in humans is the loss of a stop codon in the opioid receptor OPRM1 (6:154360569), potentially changing the structure of the protein encoded by this gene in some transcripts. Other mutations in this gene are associated to heroin addiction (Shi et al. 2002), and pain perception (Tan et al. 2009), but also sociality traits (Pearce et al. 2017). Interestingly, a recent study found a pain insensitivity disorder caused by a mutation in *ZFHX2* (Habib et al. 2017), which carries an AHMC, and three HHMCs are observed in *NPAP1*, which might be associated with the Prader-Willi syndrome, involving behavioral problems and a high pain threshold (Buiting et al. 2007). Such changes may point to differences in levels of resilience to pain between Neanderthals and modern humans.

## Conclusion

The long-term evolutionary processes that led to the human condition (Pääbo 2014) is still subject to debate and investigation, and the high-quality genomes from archaic humans provide opportunities to explore the recent evolution of our species. We want to contribute to an attempt to unveil the genetic basis of specific molecular events in the time-window after the split from these archaic populations and before the emergence of most of the present-day diversity. We sought to combine different sources of information, from genome-wide enrichment analyses to functional information available for specific genes, to identify threads linking molecular needles in this expanded haystack. In doing so, we have mainly built on existing proposals concerning brain-related changes, but we have divided the observations into different biological levels, from cellular changes through brain organization differences to complex phenotypic traits. Only future experimental work will determine which of the changes highlighted here contributed significantly to making us “fully human”. We hope that our characterization and presentation of some new candidate genes will help prioritize inquiry in this area, since the specific type of validation depends on each candidate gene or network.

## Supporting information

Supplementary Discussion

## Methods

We used the publicly available high-coverage genotypes for three archaic individuals: One Denisovan (Meyer et al. 2012), one Neanderthal from the Denisova cave in Altai mountains (Prüfer et al. 2014), and another Neanderthal from Vindija cave, Croatia (Prüfer et al. 2017). The data is publicly available under http://cdna.eva.mpg.de/neandertal/Vindija/VCF/, with the human genome version *hg19* as reference, covering ∼1.8 billion base pairs of the genome (Prüfer et al. 2017). We applied further filtering to remove sites with less than 5-fold coverage and more than 105-fold coverage in the Altai Neanderthal or 75-fold coverage in the other archaic individuals, if such cases occurred. We also removed sites with genotype quality smaller than 20, and heterozygous sites with strong allele imbalance (<0.2 minor allele frequency). Although these permissive filters increase power compared to previous studies, we caution that in some cases genotypes might be incorrect. We added the genotype and coverage for the exome and chromosome 21 sequences of the Vindija and El Sidrón Neanderthals from previous studies (Castellano et al. 2014; Kuhlwilm et al. 2016), with 75-fold and 50-fold coverage cutoffs, respectively. These studies provided data for the same Vindija individual (Prüfer et al. 2017).

We applied the Ensembl Variant Effect Predictor VEP (McLaren et al. 2016) in order to obtain inferences for protein-coding and regulatory mutations, scores for SIFT (Kumar et al. 2009), PolyPhen (Adzhubei et al. 2010), CADD (Kircher et al. 2014) and GWAVA (G.R.S. Ritchie et al. 2014), and allele frequencies in the 1000 Genomes and ExAC human variation databases (Auton et al. 2015; Lek et al. 2016). We used the inferred ancestral allele from published data on multiple genome alignments (Paten et al. 2008), and at positions where this information was not available, the macaque reference allele, *rheMac3* (*Yan et al. 2011*). We determined the allele frequencies in present-day humans using the dbSNP database build 147 (Sherry et al. 2001). We retrieved the counts for each allele type, and summarized the counts of non-reference alleles at each position. Grantham scores (Grantham 1974) were calculated for missense mutations.

Data processing and database retrieval was performed using bcftools/samtools v1.0 (Li 2011), bedtools v2.16.2 (Quinlan and Hall 2010), and R/Bioconductor (Huber et al. 2015), with rtracklayer (Lawrence et al. 2009) and biomaRt (Durinck et al. 2005) packages, and plotting with Rcircos (Zhang et al. 2013). We analyzed all positions where at least two alleles (human reference and alternative allele) were observed among the human reference and at least one out of three of the high-coverage archaic individuals, in at least one archaic chromosome. The 22 autosomal chromosomes and the X chromosome were analyzed, in the absence of Y chromosome data for the three female archaic individuals. The data for 4,409,518 segregating sites is available under [http:tbd.database]. The following subsets were created:

Fixed differences: Positions where all present-day humans carry a derived allele, while at least two out of three archaics carry the ancestral allele, accounting for potential human gene flow into Neanderthals.

High-frequency (HF) differences: Positions where more than 90% of present-day humans carry a derived allele, while at least the Denisovan and one Neanderthal carry the ancestral allele, accounting for different types of errors and bi-directional gene flow.

Extended high-frequency differences: Positions where more than 90% of present-day humans carry a derived allele, while one of the following conditions is true: a) Not all archaics have reliable genotypes, but those that have carry the ancestral allele. b) Some archaics carry an alternative genotype that is not identical to either the human or the ancestral allele. c) The Denisovan carries the ancestral allele, while one Neanderthal carries a derived allele, which allows for gene flow from humans into Neanderthals. d) The ancestral allele is missing in the EPO alignment, but the macaque reference sequence is identical to the allele in all three archaics.

We also created corresponding lists of archaic-specific changes. Fixed changes were defined as sites where the three archaics carry the derived allele, while humans carry the ancestral allele at more than 99.999%. High-frequency changes occur to less than 1% in present-day humans, while at least two archaic individuals carry the derived allele. An extended list presents high-frequency changes where the ancestral allele is unknown, but the macaque allele is identical to the present-day human allele.

A ranking of mutation density was performed for genes with protein-coding sequences and their genomic regions as retrieved from Ensembl. For each gene, unique associated changes as predicted by VEP were counted. A ranking on the number of HF changes per gene length was performed for all genes that span at least 5,000 bp in the genome and carry at least 25 segregating sites in the dataset (at any frequency in humans or in archaics), in order to remove genes which are very short or poor in mutations. The top 5% of the empirical distribution was defined as putatively enriched for changes on each lineage. The ratio of lineage-specific HF changes was calculated for the subset of genes where at least 20 lineage-specific HF changes were observed on the human and the archaic lineages combined. The top 10% of the empirical distribution was defined as putatively enriched for lineage-specific changes.

We performed enrichment tests using the R packages ABAEnrichment (Grote et al. 2016) and DescTools (Signorell 2017). We used the NHGRI-EBI GWAS Catalog (MacArthur et al. 2017), and overlapped the associated genes with protein-coding changes on the human and archaic lineages, respectively. We counted the number of HF missense changes on each lineage and the subset of those associated to each trait (“Disease trait”), and performed a significance test (G-test) against the number of genes associated to each trait, and all genes in the genome, with a P value cutoff at 0.1. This suggests a genome-wide enrichment of changes for each trait. We then performed a G-test between the numbers of HF missense changes on each lineage, and the subset of each associated to each trait (P-value cutoff at 0.1), to determine a difference between the two lineages. We then performed an empirical test by creating 1,000 random sets of genes with similar length as the genes associated to each trait, and counting the overlap to the lineage-specific missense changes. At least 90% of these 1,000 random sets were required to contain fewer missense changes than the real set of associated genes. Only traits were considered for which at least 10 associated loci were annotated.

Gene Ontology (GO) enrichment was performed using the software FUNC (Prüfer et al. 2007), with a significance cutoff of the adjusted p-value < 0.05 and a family-wise error rate < 0.05. When testing missense changes, a background set of synonymous changes on the same lineage was used for the hypergeometric test. When testing genes with relative mutation enrichment, the Wilcoxon rank test was applied. Enrichment for sequence-specific DNA-binding RNA polymerase II transcription factors and transcription factor candidate genes from (Chawla et al. 2013), and genes interacting at the centrosome-cilium interface (Gupta et al. 2015) was tested with an empirical test in which 1,000 random sets of genes were created that matched the length distributions of the genes in the test list. The same strategy was applied for genes expressed in the developing brain (Table S10) (Miller et al. 2014). Protein-protein interactions were analyzed using the STRING online interface v10.5 (Szklarczyk et al. 2017) with standard settings (medium confidence, all sources, query proteins only) as of January 2018. The overlap with selective sweep screens considers HHMCs within 50,000 bp of the selected regions (Prüfer et al. 2014; Zhou et al. 2015; Peyrégne et al. 2017).

## Acknowledgments

We thank S. Han and T. Marques-Bonet for helpful discussions, and A. G. Andirkó and P .T. Martins for help with figures. M.K. is supported by a Deutsche Forschungsgemeinschaft (DFG) fellowship (KU 3467/1-1). C.B. acknowledges research funds from the Spanish Ministry of Economy and Competitiveness/FEDER (grant FFI2016-78034-C2-1-P), Marie Curie International Reintegration Grant from the European Union (PIRG-GA-2009-256413), research funds from the Fundació Bosch i Gimpera, MEXT/JSPS Grant-in-Aid for Scientific Research on Innovative Areas 4903 (Evolinguistics: JP17H06379), and Generalitat de Catalunya (Government of Catalonia) – 2017-SGR-341.

## Author contributions

M.K. and C.B. analyzed data and wrote the manuscript.

