## Supplementary Discussion for "A catalog of single nucleotide changes distinguishing modern humans from archaic hominins"

**This PDF includes:**

Supplementary Discussion

Supplementary Table and Figure legends

Supplementary References

#### Supplementary discussion

### 1. Cell division

Among the 15 fixed protein-coding changes identified here but absent from previous analyses (Pääbo 2014; Prüfer et al. 2014), some might also contribute to complex modifications of pathways in cell division: The AHR protein is involved in cell cycle regulation (Puga et al. 2002) and shows an excess of HF changes on the human lineage, the dynein DNHD1 might be recruited to the kinetochore (Bader et al. 2011) and is overexpressed in fetal brain (Safran et al. 2010), and the SSH2 protein (two fixed changes, one of which is first described here; and one on the archaic lineage) might interact with spindle assembly checkpoint proteins (Bailey et al. 2015). SHROOM4, which is associated to a mental retardation syndrome with delayed speech and aggressive behavior (Stocco dos Santos et al. 2003), may also be relevant (Yoder and Hildebrand 2007). Other proteins that carry two HHMCs are involved in mitosis, for example the spindle checkpoint regulator CHEK1 (Zachos et al. 2017), the Dynein Axonemal Heavy Chain 1 (encoded by *DNAH1*), the mitotic regulator AZI1 (*CEP131*) (St-Denis et al. 2017), the Cyclin D2 (*CCND2*) and the Protein Tyrosine Phosphatase Receptor Type C (*PTPRC*) (Melkerson-Watson et al. 1994). Other genes with HHMCs that could be part of the same functional network are *KIF26B* (Wojcik et al. 2018), *DCHS1* (Cappello et al. 2013), *FOXM1* (Thiru et al. 2014) and *FMR1* (Callan et al. 2010), which carry a putative enrichment of HF changes, and *TOP2A* (Yoshida and Azuma 2016). The TOP2A protein shows the largest number of interactions (53) with other HHMC-carrying proteins, while CHEK1, KIF18A, KIF15 and PTPRC are among highly-interacting proteins with more than ten interactions, which suggests that these proteins might function as interaction hubs in modifications of the cell division complex. Furthermore, enrichment in cell-cycle related GO categories has been found for candidate regions for ancient positive selection (Racimo 2016), and *ANAPC10* has been highlighted, containing two potentially disruptive intronic changes that are fixed derived in modern humans and ancestral in both Neanderthals and Denisovans. This gene carries a total of 39 HF changes (11 of them fixed) specific to modern humans, but none for archaics.

On the archaic lineage, we find an AHMC in the *ASPM* gene, along with 24 other HF changes, but none in modern humans, resulting in an excess of archaic SNCs. The proteins ASPM and CIT, which carries an AHMC that is listed among the most disruptive non-synonymous derived SNCs in archaics (Table S31 in (Castellano et al. 2014)), are known to co-localize to the midbody ring during cytokinesis and regulate spindle orientation by affecting the dynamics of astral microtubules (Gai et al. 2016). These proteins regulate astral microtubules and thus the orientation of cell division in archaics, whereas in modern humans we find proteins regulating kinetochore microtubules, thus the timing of cell division. This difference could indicate two alternative ways of modulating cell division on the different lineages.

### 2. Cellular properties of neurons

The establishment of new connections requires protection, particularly as some of these connections reach long distance and are associated with enhanced activity following rewiring events, like for vocal motor neurons in songbirds (Pfenning et al. 2014). The gene *MAL*, which is implicated in myelin biogenesis and function, shows up in selective sweep regions (Racimo 2016; Peyrégne et al. 2017) and is enriched for HF changes on the human lineage, while its orthologue *MAL2* carries a HHMC. A gene with HHMCs that is associated with the organization of the axon initial segment and nodes of Ranvier during early development is *NFASC* (Ango et al. 2004). The protein encoded by this gene is a L1 family immunoglobulin cell adhesion molecule, and we find that also the *L1CAM* gene carries an AHMC (Pollerberg et al. 2013). NFASC is also an interactor of DCX (Yap et al. 2012), which might have been under positive selection in humans (Peyrégne et al. 2017) and is enriched for HF SNCs on the human lineage, but carries an AHMC as well. At least two genes associated with the process and timing of myelination, *PTEN* (Harrington et al. 2010), *VCAN* (Dours-Zimmermann et al. 2009) and *NCMAP* (Ryu et al. 2008) are among genes with an excess of HF SNCs in modern humans. Other genes carrying HHMCs in our dataset associated with myelination include *SCAP* (Verheijen et al. 2009), *RB1CC1* (Menzies et al. 2015), *TENM4* (Hor et al. 2015), *CDKL1* (Hsu et al. 2011) and *ADSL* (Jurecka et al. 2012), and genes with an excess of changes on the human lineage with similar functions include *FBXW7* (Kearns et al. 2015), *KIFAP3* (Morfini et al. 2009), and *AMPH* (Butler et al. 1997). The AMPH protein interacts closely with the huntingtin protein HTT (which also carries a HHMC) and is involved in myelination processes (Huang et al. 2015).

Another interesting class that emerges from the set of genes is related to synaptic vesicle endocytosis, critical to sustain a high rate of synaptic transmission. We find a formal enrichment of genes with an excess of HF changes on the human compared to the archaic lineage with gene products located in the postsynaptic membrane and dendrites. *PACSIN1* (Widagdo et al. 2016) carries a HHMC, is among genes with an excess of HF changes, and has been highlighted as putatively under positive selection on the human lineage, along with other synaptic plasticity related genes such as *SIPA1L1* (Zhou et al. 2015; Racimo 2016; Peyrégne et al. 2017), *SH3GL2* (Arranz et al. 2015) and *STX1A* (Craig et al. 2015). Among genes harboring HHMCs and related to synaptic vesicle endocytosis, we find *LMNB2* (Razafsky et al. 2016) and *SV2C* (Janz and Südhof 1999). Finally, *SYT1*, which is critical for synaptic vesicle formation (Lee and Littleton 2015), carries a deleterious HHMC (Table 2).

Synaptic properties have been mentioned before in the context of human specific traits, for instance in postnatal brain development in humans, chimpanzees and macaques (Liu et al. 2012), with a focus on synaptogenesis and synaptic elimination in the prefrontal cortex. A period of high synaptic plasticity in humans has been related to a cluster of genes around a transcription factor encoded by the *MEF2A* gene. Even though this gene neither carries a protein-altering change nor shows a particular pattern in our analysis, any of the 26 HF SNCs it harbors on the modern human lineage could have had a functional impact not captured here. Apart from that, several of the genes with an excess of HF changes in modern humans do belong to this cluster: *CLSTN1, FBXW7, GABBR2, NRXN3, PTPRJ, PTPRN2, SLIT3*, and *STX1A*, three of which (*CLSTN1, FBXW7* and *STX1A*) are associated with signals of positive selection (Peyrégne et al. 2017). In addition, the above-mentioned AMPH interacts via CDKL5 (Sekiguchi et al. 2013) with HDAC4 (Trazzi et al. 2016). The latter exhibits an excess of HF changes in modern humans, and is known to repress the transcriptional activation of *MEF2A* (Miska et al. 1999). A putative signature of positive selection upstream of *MEF2A* (Somel et al. 2014) suggests that this may be part of a broader network which might be supported by our analysis. Finally, *ENTHD1/CACNA1I*, which contains a HHMC that can no longer be considered as fixed, but occurs at a very high frequency (>99.9%), lies in a selective sweep region (Peyrégne et al. 2017). The protein encoded by this gene is involved in synaptic vesicle endocytosis at nerve terminals (Ryan 2006) and is regulated by the *MEF2* gene family (Kornilov et al. 2016).

### 3. The brain growth trajectory

Changes in genes that influence microcephaly are found on both lineages: In archaics there are AHMCs in the microcephaly candidate genes *ASPM* (Tungadi et al. 2017) and *CIT* (Bianchi et al. 2017). The ASPM-katanin complex controls microtubule disassembly at spindle poles and misregulation of this process can lead to microcephaly (Jiang et al. 2017), which is of interest given the presence of a HHMC in *KATNA1* and a fixed non-coding change in *KATNB1*, while no such changes were observed in archaics (Yigit et al. 2016). Other genes associated with microcephaly that harbor non-synonymous SNCs are *CASC5* (two in humans, one in archaics) (Genin et al. 2012), *CDK5RAP2* (in humans), *MCPH1* (in archaics) (Arroyo et al. 2017), *ATRX* (one in humans and archaics each) (Ritchie et al. 2014), and *NHEJ1* (El Waly et al. 2015) (a deleterious one in humans, and one in archaics). Disease mutations in *SCAP* or *ADSL* have also been associated with microcephaly phenotypes as well (Suzuki et al. 2013; Jurecka et al. 2015), and Formin-2 (*FMN2*), which carries a deleterious regulatory change in modern humans, influences the development of the brain causing microcephaly in mice (Lian et al. 2016).

The SPAG5 protein, which carries three fixed HHMCs, has been claimed to interact with CDK5RAP2 (Kodani et al. 2015), is a direct target of PAX6 (Asami et al. 2011), via which it affects cell division orientation, and therefore is critical in the course of brain development. The *SPAG5* gene might be a particular example of specific consequences for a relevant SNC on the human lineage: One of the three fixed non-synonymous changes in the SPAG5 protein is a Proline-to-Serine substitution at position 43. This position is phosphorylated in humans (Dephoure et al. 2008) during the mitotic phase of the cell cycle, directly through the protein phosphatase 6 (PPP6C) at the Serine at this position (Rusin et al. 2015), with the effect of a modification of the duration of the metaphase. PPP6C regulates the mitotic spindle formation (Zeng et al. 2010), and the *PPP6C* gene itself carries five HF SNCs on the modern human lineage, one of which is a transcription factor binding site (for HNF4A/HNF4G), and only one SNC on the archaic lineage. This specific substitution in *SPAG5* seems likely to influence the duration of the metaphase through phosphorylation, as a molecular consequence of this HHMC.

Among macrocephaly-related genes with HHMCs in *RNF135* (Douglas et al. 2007)*, CUL4B* (Tarpey et al. 2007) and *CCND2* (Mirzaa et al. 2014), the latter also shows a large number of HF changes on the human lineage, and the HHMC in *CUL4B* is inferred to be deleterious (Table 2). Other macrocephaly candidates such as *NFIX* (Klaassens et al. 2014)*, NSD1* (Buxbaum et al. 2007) and *GLI3* (Jamsheer et al. 2012) have been claimed to have played an important role in shaping the distinctly modern human head (Gokhman et al. 2017) and show numerous SNCs in non-coding regions. *GLI3* might have been under positive selection (Peyrégne et al. 2017) and carries 20 HF SNCs on the human, but only one on the archaic lineage. Two of the very few genes hypothesized to regulate expansion and folding of the mammalian cerebral cortex by controlling radial glial cell number and fate, *TRNP1* (Stahl et al. 2013) and *TMEM14B* (Liu et al. 2017), exhibit HF 3'-UTR changes in modern humans, and *TRNP1* shows an excess of changes on the modern human lineage. The expression of these two genes in the outer subventricular zone might be important (Martínez-Martínez et al. 2016), since this is an important region for complexification of neocortical growth in primates (Dehay et al. 2015), and for which an enriched activation of mTOR signaling has been reported (Nowakowski et al. 2017). In addition to other genes in the mTOR-pathway, such as *PTEN* (Li et al. 2017) or *CCND2*, two possibly interacting modulators (Cloutier et al. 2017) of the mTOR signaling pathway stand out in our dataset: *ZNHIT2* with one deleterious SNC (Table 2) might have been under positive selection (Peyrégne et al. 2017), and *CCT6B* carries a deleterious change according to both SIFT and C-score. The transcription factor encoded by *RB1CC1* is essential for maintaining adult neuronal stem cells in the subventricular zone of the cerebral cortex (Wang et al. 2013). This gene carries a HHMC, a regulatory SNC that has been suggested to modify transcriptional activity (Weyer and Pääbo 2016), and a signature of positive selection (Prüfer et al. 2014).

The number of HHMCs that putatively interact with proteins at the centrosome-cilium interface (Gupta et al. 2015) is more than expected using 1,000 random gene sets of a similar length distribution, for which 98.9% contain fewer genes with HHMCs. However, 99.9% of random sets also contain fewer genes with AHMCs, suggesting that differences between humans and archaics might lie in the particular genes rather than their numbers. On the archaic side, an enrichment of genes with AHMCs associated to “Corneal structure” may relate to archaic-specific changes in brain growth-trajectories since the size and position of the frontal and temporal lobes might affect eye and orbit morphology (Pereira-Pedro et al. 2017), and the macrocephaly-associated gene *RIN2* (Basel-Vanagaite et al. 2009) carries an AHMC.

### 4. The impact on cognition

It has long been hypothesized that language and its neurological foundation were important for the evolution of humans and uniquely human traits, closely related to hypotheses on the evolution of cognition and behavior. It is noteworthy that among traits associated with cognitive functions such as language or theory of mind, the timing of myelination appears to be a good predictor of computational abilities (Skeide and Friederici 2016; Grosse Wiesmann et al. 2017). We suggest that some genes with changes on the human lineage might have contributed more specifically to cognition-related changes, although we admit that the contribution of single SNCs to these functions is less straightforward than their contribution to molecular mechanisms, since disease mutations in many genes may have disruptive effects on cognitive abilities. The basal ganglia are a brain region where *FOXP2* expression is critical for the establishment and maintenance of language-related functions (Vargha-Khadem et al. 2005; Enard et al. 2009), and several genes carrying HHMCs have been described previously as important for basal ganglia functions. The HTT protein has long been implicated in the development of Huntington’s disease, which is associated with corticostriatal dysfunction, and is known to interact with FOXP2 (Hachigian et al. 2017). Mutations in *SLITRK1,* which might have been under positive selection (Peyrégne et al. 2017), have been linked to Tourette's syndrome, a disorder characterized by vocal and motor tics, resulting from a dysfunction in the corticostriatal-thalamocortical circuits (Abelson et al. 2005). NOVA1 regulates RNA splicing and metabolism in a specific subset of developing neurons, particularly in the striatum (Jelen et al. 2010). As pointed out above, *NOVA1* is an interactor of *ELAVL4*, which belongs to a family of genes known to promote the production of deep layer *FOXP2-*expressing neurons (Konopka et al. 2012; Alsiö et al. 2013; Popovitchenko et al. 2016), and part of a neural network-related cluster that has been highlighted as putatively under positive selection in humans (Zhou et al. 2015). Within this network, α-synuclein (encoded by *SCNA*) might serve as a hub and is specifically expressed in brain regions important for vocal learning regions in songbirds (Pfenning et al. 2014). *SCNA* and *SV2C*, which carries a HHMC, are involved in the regulation of dopamine release, with *SV2C* expression being disrupted in *SCNA*-deficient mice and in humans with Parkinson’s disease (Dunn et al. 2017). Genes in the cluster of selection signals (Zhou et al. 2015) are implicated in the pathogenesis of Alzheimer’s disease, which (together with Huntingon’s and Parkinson’s diseases) is linked to a *FOXP2*-driven network (Oswald et al. 2017). Some introgressed archaic alleles are downregulated in specific brain regions (McCoy et al. 2017), especially pronounced in the cerebellum and basal ganglia. One notable example is *NTRK2*, which shows an excess of HF changes on the human lineage and a signature of positive selection (Peyrégne et al. 2017), and is also a FOXP2 target (Vernes et al. 2011), a connection which has been highlighted for the vocal learning circuit in birds (Hilliard et al. 2012). Other genes harboring HHMCs such as *ENTHD1* (Kornilov et al. 2016) and *STARD9* (Chen et al. 2017), as well as genes in introgression deserts (Vernot et al. 2016), have been associated with language deficits. It may indeed have taken a complex composite of changes to make our brain fully language-ready (Boeckx and Benítez-Burraco 2014), where not all changes needed to reach fixation.

Two genes linked to Alzheimer’s are *PTEN* (Ferrarelli 2016), and *RB1CC1* (Chano et al. 2007). Among genes with deleterious HHMCs, *SLC6A15* has been associated to emotional processing in the brain (Choi et al. 2016), and may be part of modifications in glutamatergic transmission (Santarelli et al. 2016), a category found in selective sweep regions (Theofanopoulou et al. 2017). *GPR153*, which carries one HHMC and two AHMCs, influences behavioral traits like decision making in rats, and is associated with various neuropsychiatric disorders in humans (Sreedharan et al. 2011). For the Adenylosuccinate Lyase (*ADSL*) the ancestral Neanderthal-like allele has not been observed in 1,000s of modern human genomes. This gene has been associated to autism (Fon et al. 1995), is part of behavioral traits like “aggressive behavior” which have been found to be enriched on the human lineage (Castellano et al. 2014), and several studies detected a signal of positive selection in modern humans (Racimo et al. 2014; Racimo 2016; Peyrégne et al. 2017). These observations make *ADSL* a strong candidate for human-specific features, particularly in light of the fact that the relevant HHMC is located in a region that is highly conserved and lies close to the most common disease mutation leading to severe adenylosuccinase deficiency (Racimo 2016). Other relevant genes, similar to *ADSL* in carrying a fixed HHMC and being frequently found in selective sweep screens, are *NCOA6*, which might be related to autism as well (Takata et al. 2018), and *SCAP.* Downregulation of the cholesterol sensor encoded by this gene has been shown to cause microcephaly, impaired synaptic transmission and altered cognitive function in mice (Suzuki et al. 2013). We want to emphasize that the networks presented in the previous sections influencing brain growth and neural wiring are likely to impact cognitive functions, since disruptions in these networks would impair the healthy human brain. Furthermore, we find an enrichment of AHMCs in genes associated to Parkinson’s disease and “Attention deficit hyperactivity disorder and conduct disorder”, suggesting that changes may have taken place in related networks on the archaic lineage as well.

#### Supplementary Table and Figure legends

**Table S1**: List of HHMCs and genomic features

**Table S2**: List of low-confidence HHMCs and genomic features

**Table S3**: List of AHMCs and genomic features

**Table S4**: List of low-confidence AHMCs and genomic features

**Table S5**: Top 5% of genes by HF SNC density on the modern human and archaic lineages, and top 10% of genes by relative excess of HF SNCs on one lineage over the other.

**Table S6**: GO enrichment for genes with relative excess of HF SNCs on the human over the archaic lineage

**Table S7**: GWAS enrichment for genes with HHMCs or AHMCs

**Table S8**: Number of interactions among genes with HHMCs or AHMCs

**Table S9**: Genes with HHMCs that are transcription factors, or at the centrosome interface

**Table S10**: Enrichment in developing brain zones for genes with HHMCs or AHMCs, proportion of random gene sets with larger overlap (Methods).

**Figure S1**: Distribution of missense and non-synonymous HF SNCs across chromosomes.

**Figure S2**: STRING graph of interactions among genes with HHMCs.
